# How Visual Context Influences Lateral Stepping Regulation While Walking on Winding Paths

**DOI:** 10.64898/2026.01.17.700122

**Authors:** Anna C. Render, Tarkeshwar Singh, Joseph P. Cusumano, Jonathan B. Dingwell

**Affiliations:** Department of Kinesiology, The Pennsylvania State University, University Park, PA, United States; Department of Engineering Science & Mechanics, The Pennsylvania State University, University Park, PA, United States

## Abstract

Goal-directed walking involves regulating foot placements to achieve specific tasks. This requires visuomotor integration. Perceptual, cognitive, and contextual salience guide attention and motor planning for navigation. Here, we quantified how perceptual salience informs lateral foot placement while walking. Participants walked along prescribed virtual paths (straight or winding), thus keeping contextual salience (the task itself) constant. We manipulated perceptual salience by systematically altering environment richness (rich vs. sparse) and path color contrast (high vs. low). We thus tested the extent to which stepping control is determined only by the walking path (i.e. its shape), or is also influenced by task-relevant salience of the paths (only), and/or the salience of the surrounding environment. We quantified head pitch angle to approximate gaze direction. We quantified lateral stepping regulation from a Goal Equivalent Manifold framework. On winding (vs. straight) paths, participants looked down more (lower mean head pitch) and more consistently (less variable head pitch). They took narrower, more variable steps and corrected stepping errors more strongly. This confirmed the predominant influence of task (contextual salience) on stepping control. On low (vs. high) contrast paths, participants looked down more and exhibited greater variability in lateral position on their path with weaker error correction. Higher contrast paths elicited stronger and more consistent stepping control. Walking in sparse (vs. rich) environments yielded somewhat less consistent, but still significant changes in head pitch and stepping regulation. Salience manipulations affected stepping differently on straight versus winding paths. Therefore, perceptual salience influences step-to-step control during naturalistic walking.

## INTRODUCTION

Walking is among the most common natural behaviors in which humans engage. Natural walking is *goal-directed*: i.e., we walk to reach a particular destination (1) or navigate fixed features of our terrain (2, 3), and to avoid obstacles (4) and/or other people (5, 6) along the way. In such continually-evolving contexts, walking and navigating our world intrinsically involves multiple aspects of embodied decision-making (7–11). People can quickly adjust their movements in real time during ongoing reaching tasks (12, 13). But they can also make similarly fast stepping adjustments while walking (14). Importantly, the biomechanics (dynamics) of the task being performed directly shape movement decisions people make during ongoing actions, for both reaching movements (15) and during walking (16, 17).

The goal-directed nature of walking involves regulating our foot placements as we walk to achieve specific desired tasks (18, 19). This goal-directed walking is achieved through distributed neural networks that coordinate voluntary movement with automatic postural control. The cerebral cortex, particularly the posterior parietal and premotor areas, plays a crucial role in step-by-step regulation of foot placement to achieve specific locomotor goals (20, 21).

In particular, walking relies heavily on visuomotor integration (22–25) and active sensing (26, 27). Vision is particularly crucial for navigation (5, 23, 28, 29) and locomotor control, acting through subcortical and cortical regions (30). The dorsal visual stream provides spatial information for online control of locomotion. The superior colliculus and pulvinar nucleus coordinate with cortical areas to support anticipatory gaze behaviors to enable proactive motor planning (31, 32). Hence, the nervous system can integrate visual information and engage predictive control mechanisms to respond to upcoming challenges (3, 33, 34). In particular, central vision allows walkers to use optic flow to guide walking (35), whereas peripheral vision seems particularly important for monitoring features of the environment (35, 36) needed to enact online stepping adjustments (22). Importantly, during natural walking, multiple factors compete for gaze attention (37, 38). Humans thus use gaze to reduce uncertainty of *task-relevant* visual information (23, 24, 28, 38, 39) to inform motor planning (21). Various aspects of visual processing likely contribute to these processes. For instance, increased color contrast enhances pupillary responses (40, 41) and attention-related neural signatures (42). However, there remains a critical gap in understanding how these processes influence real-world locomotor behavior.

A key concept in understanding how humans use visual information to navigate their environments as they walk is that of visual *salience* (43–45), which refers to the degree to which certain aspects of a visual scene “stand out” visually and thus draw one’s visual attention. Caduff and Timpf (46) proposed a conceptual framework that distinguishes three key neural processes: Perceptual salience, Cognitive salience, and Contextual salience. *Perceptual Salience*, a bottom-up sensory process, is driven by visual properties of the environment that readily capture attention, like color, size, or shape. *Cognitive Salience*, top-down attentional control, is influenced by prior knowledge and memory; it directs attention to features relevant to navigation tasks through mental representations. *Contextual Salience*, task dependent modulation, is affected by the navigation task’s purpose and mode of transportation. It modulates attention by allocating resources based on task demands. Perceptual Salience engages early visual areas and the ventral attention network. Cognitive salience recruits prefrontal and dorsal visuomotor regions. Contextual salience involves goal-directed attention networks to identify potential landmarks (46). Together, these elements illustrate the interaction between environmental influences and cognitive processes, emphasizing that effective navigation depends on the flexible integration of sensory signals and task objectives.

Here, we examined how variations in visual input, specifically perceptual salience, affect the step-to-step control of walking during path following. Participants (healthy adults) walked along either of two specified paths (straight or winding). We kept the path widths and shapes the same, ensuring consistent navigation goals and therefore constant contextual salience for each path. We then manipulated perceptual salience by independently varying the richness of the peripheral environment (e.g., rich forest versus sparse plain) and the walking path’s visual contrast (e.g., high versus low contrast). Cognitive salience acts as a bridge between the unchanging contextual task and the changing perceptual stimuli. It is in this space that participants process and integrate these perceptual changes into their navigation strategies. By keeping contextual salience constant while systematically manipulating perceptual salience, our experimental design isolated how variations in bottom-up visual processing influence motor output through cognitive integration networks. This integration guides participants’ navigation strategies and, ultimately, the action component of the perception-action cycle, allowing for adaptive walking behaviors in response to environmental changes.

To assess how manipulating these visual processes affected locomotor control, we assessed stepping regulation (18, 19, 47, 48) using a Goal Equivalent Manifolds (GEMs) framework (49). People regulate their foot placements from step to step to achieve specific task goals (48), such as following defined (or stereotyped) paths (50). When walking on straight paths, regulating lateral stepping involves a dynamic trade-off between maintaining step width and maintaining lateral position relative to the path (19). As such, egocentric information (i.e., of where the walker is within its environment) derived from vision (21, 22, 29) provides the contextual salience needed for such regulation: i.e., to achieve *goal-directed* walking, one must first know where the (external) “goals” are (18). Humans readily adapt their lateral stepping to changing environmental conditions, supporting the notion that humans make “embodied decisions” (8, 51) regarding how to regulate their stepping movements in real-time during ongoing locomotion. Humans systematically alter their lateral stepping to navigate slowly narrowing (52), or winding pathways (47), or to perform lateral maneuvers (53). Such skills are essential for addressing unforeseen obstacles or shifts in terrain. Our prior work thus demonstrated how manipulating task goals (i.e., contextual salience) affects walking control. This present study extends those ideas to quantify how perceptual and cognitive salience contribute to stepping regulation.

Our overall guiding hypothesis is that walking movements arise from continuous decision-making action-perception processes (9, 22) – i.e., each step taken affects the decision of where to step next. Our previous computational models of stepping regulation (19, 48, 53) presumed stepping behaviors are guided solely by *contextual* salience (i.e., the walking task itself) and therefore that the visual input of the task is (a least) “sufficient” for this task. Further, adding stepping regulation to a dynamic walking model (18) confirmed that indeed, *external* state information (i.e., where the body is in the world), in addition to knowledge *internal* (body) states, was necessary for the model to enact goal-directed walking behaviors like those exhibited by humans (18, 19). In humans, this external state information would typically be acquired through vision (21, 22, 29). Our prior experiments that manipulated contextual salience supported our stepping model predictions (47, 52, 53). In the present experiment, the *task* was to walk on the path presented, and all paths were visible. Since the task requirements (i.e., walk on the path) remained unchanged, there was no biomechanical or physiological requirement that individuals alter their stepping behavior. Thus, one real plausible outcome was that neither of the imposed manipulations of visual input (i.e., either of the peripheral environment or of the path) would induce participants to change their stepping behavior. This would be expected if only contextual salience were relevant to stepping control and if being able to see one’s environment and walking path are indeed sufficient to provide that contextual salience.

However, multiple studies of gaze during walking support the idea that gaze is used primarily to enhance *task*-relevant visual information (3, 38) and reduce visual uncertainty (23, 24, 45). If so, one alternative plausible outcome would be to expect that decreasing the perceptual salience (i.e., visual contrast) of the paths themselves would degrade task-relevant visual information and thus degrade participants’ stepping control, whereas altering the perceptual salience of the surrounding peripheral environment (which is *not* task-relevant) would have little-to-no effect on stepping regulation.

However, multiple prior studies have also highlighted the potential importance of peripheral vision in governing locomotor behavior (22, 35, 36, 54). Hence, a third alternative plausible outcome would be to expect that decreasing the perceptual salience of either the peripheral environment or the paths themselves (or both) would induce participants to change their stepping behavior.

The experimental design adopted here employed a naturalistic walking task while holding contextual salience constant for each walking path (straight or winding) presented. By then manipulating the perceptual salience of the peripheral environment and walking paths independently, we intended to directly identify the extent to which both central and peripheral visual perceptual salience influence motor planning of stepping through visuomotor integration networks. Here, as the geometric characteristics of each path remained unchanged, any changes in stepping behavior would occur independently of each task’s biomechanical demands, and instead arise as adaptive responses to the imposed variations in the salience of the available visual information.

## MATERIALS AND METHODS

### Participants

Prior to participation, all participants provided written informed consent, as approved by the Institutional Review Board of The Pennsylvania State University. Twenty-eight young healthy adults (Table 1) participated. Participants were screened to ensure no medications, lower limb injuries, surgeries, musculoskeletal, cardiovascular, neurological, or visual conditions affected their gait.

**Table 1.**
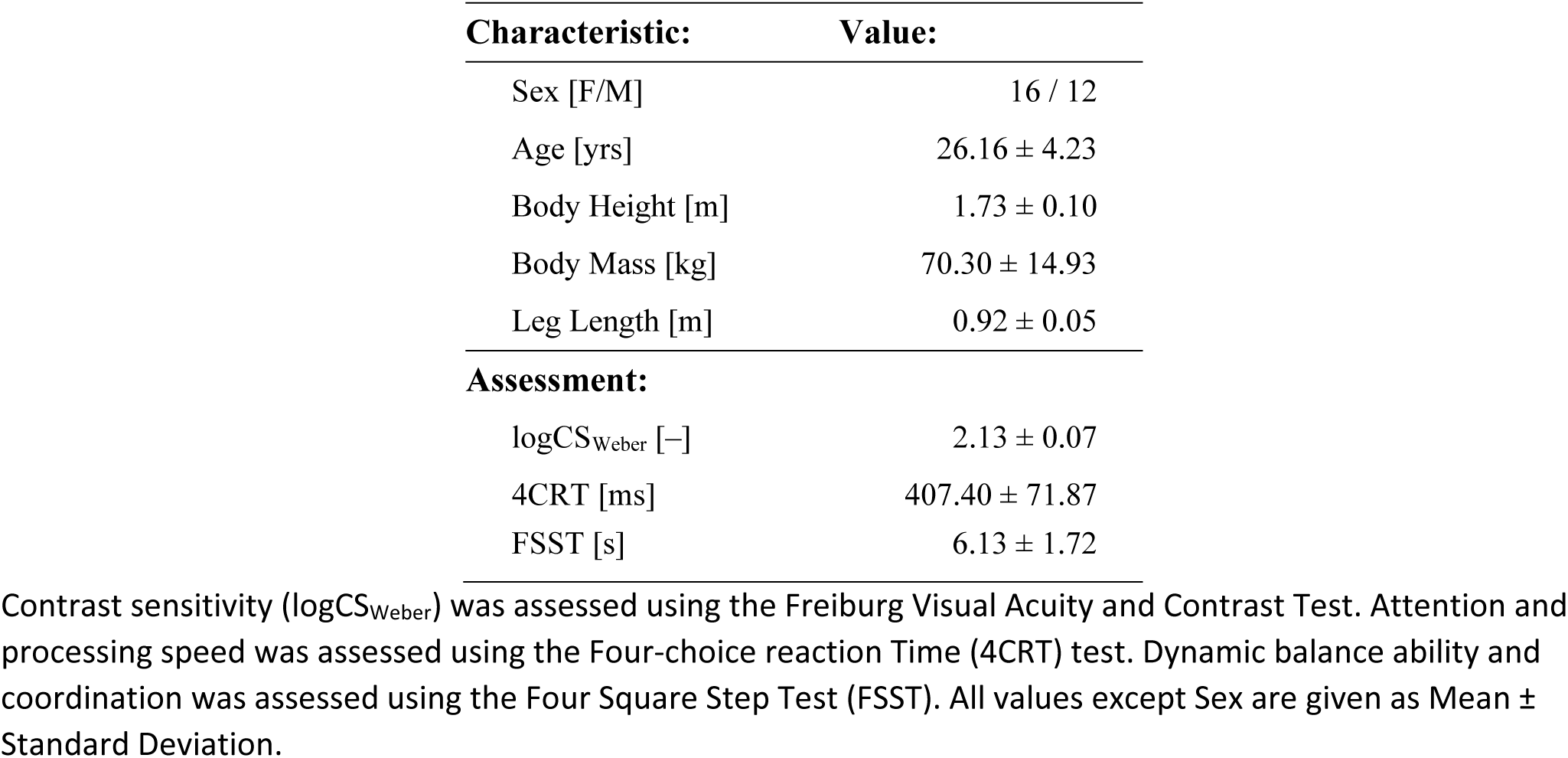
Participant physical characteristics and assessment scores.

| Characteristic: | Value: |
| --- | --- |
| Sex [F/M] | 16 / 12 |
| Age [yrs] | 26.16 ± 4.23 |
| Body Height [m] | 1.73 ± 0.10 |
| Body Mass [kg] | 70.30 ± 14.93 |
| Leg Length [m] | 0.92 ± 0.05 |
| **Assessment:** |  |
| logCS<sub>Weber</sub> [–] | 2.13 ± 0.07 |
| 4CRT [ms] | 407.40 ± 71.87 |
| FSST [s] | 6.13 ± 1.72 |

Contrast sensitivity (logCS<sub>Weber</sub>) was assessed using the Freiburg Visual Acuity and Contrast Test. Attention and processing speed was assessed using the Four-choice reaction Time (4CRT) test. Dynamic balance ability and coordination was assessed using the Four Square Step Test (FSST). All values except Sex are given as Mean ± Standard Deviation.

### Assessments

We measured participants’ visual contrast sensitivity using the Freiburg Visual Acuity and Contrast Test (55), administered using the FrACT_10_ on-line web application (version 1.5, release date 2023; https://michaelbach.de/fract/). The FrACT_10_ contrast test presents participants with Landolt-C’s that vary over a wide range of visual contrasts (32 trials) and asks them to identify correct responses using an 8-alternative forced choice procedure (55). Weber contrast thresholds (C_Weber_) are then converted to logarithmic Weber contrast sensitivity scores: logCS_Weber_ = log_10_(1/C_Weber_) (56, 57) (Table 1).

We assessed participants’ attention and processing speed using the Four-Choice Response Task (4CRT; Table 4-1) (58), administered using PEBL software (https://pebl.sourceforge.net/) (59). Each participant completed two blocks of 50 stimuli and we recorded their mean reaction time.

We evaluated participants’ balance using the Four Square Step Test (FSST; Table 4-1) (60). Each participant completed three trials and we recorded their mean time.

### Apparatus and Stimuli

Participants walked in a M-Gait virtual reality system (Motek Medical, Netherlands; Fig. 1A) equipped with a 1.2 m wide treadmill. Each participant wore a safety harness secured to the ceiling overhead. Such high-fidelity virtual environments that incorporate naturalistic movement control can yield gaze behavior similar to walking in the real world (61).

**Figure 1.**
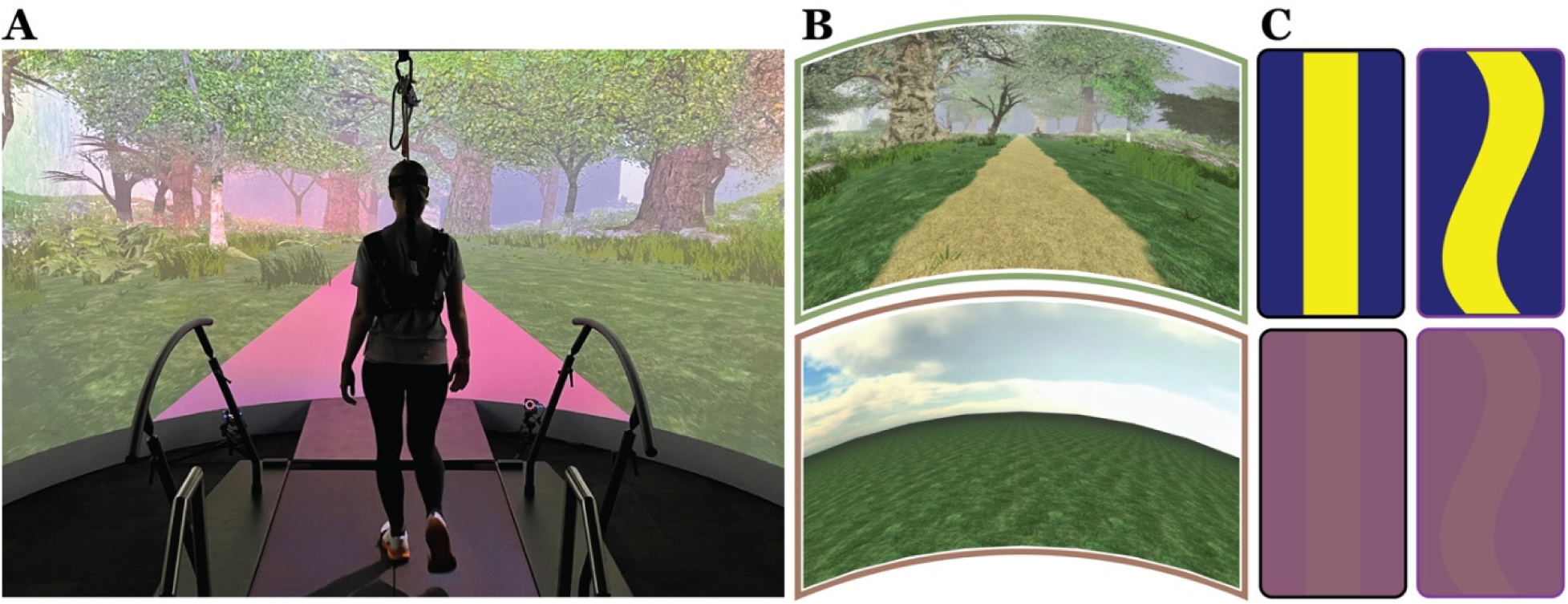
Experimental Setup and Test Conditions. **A)** Photo of a participant walking in the Motek M-Gait virtual reality (VR) system. **B)** Participants walked through each of two virtual environments: a visually rich forest scene with dense foliage (“Rich”; top) and a flat open grassy plain (“Sparse”; bottom). **(C)** In each environment, each participant walked on paths that were either Straight (STR; left) or Winding (HIF; right), and of either High (HC; top) or Low (LC; bottom) visual color contrast.

#### VR Environments

The Rich environment featured a forest scene with dense foliage (Fig. 1B; top), while the Sparse environment was simply a flat open grassy plain (Fig. 1B; bottom).

#### Path Contrast

We selected color combinations for the path and its background to create both high and low contrast combinations using spherical CIELAB color space (41). We chose colors along the blue/yellow axis.

We computed net luminance (*L_Net_*) of the combined path and background based on their relative widths using the luminance (L) values from the CIELAB [L,a,b] color coordinates. All paths were 0.45 m wide on a 1.2 m wide background therefore, the *L_Net_* was:

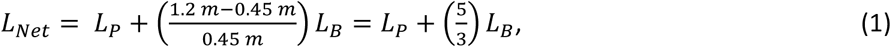

where *L_P_* is the luminance of the path, and *L_B_* is the luminance of the background. We held net luminance constant between the high and low contrast combinations. High contrast (HC) conditions projected a yellow path [L,a,b] = [97.0,-22.0,95.0]; corresponding to [R,G,B] = [254,255,0], onto a dark blue background [L,a,b] = [13.0,47.5,-65.0]; [R,G,B] = [0,0,129] (Fig. 1C; top). This yielded a contrast ratio of *L_P_/L_B_* = 7.46 for the HC paths.

To determine the color combination for the low contrast (LC) conditions, we linearly interpolated between the HC blue background and the HC yellow path, shifting each proportionally towards the other in CIELAB space. Larger color shifts reduce the difference between the resulting colors, making the path and background colors harder to visually discriminate. We shifted the blue background 36% of the linear distance toward the yellow path color:

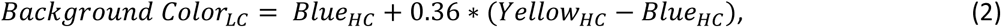

resulting in a LC background color of [L,a,b] = [43.24,22.48,-7.4]; [R,G,B] = [134,88,115]. To maintain constant net luminance (*L_Net_*), the path color was shifted proportionally as:

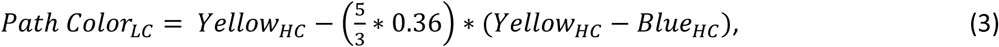

resulting in a LC path color of [L,a,b] = [46.6,19.7,-1.0]; [R,G,B] = [141,98,113] (Fig. 1C; bottom). This yielded a contrast ratio of *L_P_/L_B_* = 1.08 for the LC paths, an almost 7-fold reduction in path contrast ratio compared to the HC condition (Fig. 1C).

#### Path Shapes

We tasked participants to walk on either straight (STR) or winding (HIF) paths projected onto the treadmill belt. We designed the winding path to oscillate pseudo-randomly as (47, 62):

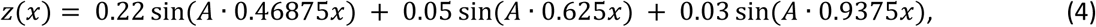

where z is the lateral position (in meters) of the path center, A is a frequency scaling factor, and x is forward treadmill distance (in meters). The oscillation frequency for the HIF path was set to A = 4, as in (47, 62).

For each, we instructed participants to “stay on the path” and encouraged them to minimize stepping errors. They received visual (firework) and auditory (flute) penalties when steps landed outside the path boundaries.

### Experimental Protocol

On each path (STR & HIF), participants walked with each combination of peripheral environment (Rich & Sparse) and path contrast (HC & LC), yielding 8 total conditions (2×2×2). For all walking trials, treadmill speed was set to 1.2 m/s and optic flow speed was matched to the treadmill speed.

Participants initially walked 3-minutes to acclimate to the system. For each condition, participants completed a 1-minute introductory trial, followed by two 3-minute experimental trials. Conditions were presented in pseudo-random order to each participant, such that order-of-presentation between conditions was counterbalanced across participants using a Latin Square design to minimize order effects. Participants were allowed to rest as needed after each trial.

### Data Acquisition and Processing

Each participant wore 17 retroreflective markers: five around the head, four around the pelvis (left/ right PSIS and ASIS), and four on each of the left and right shoes (aligned with first and fifth metatarsal heads, lateral malleolus, and calcaneus). Kinematic data were collected at 100 Hz from a 10-camera Vicon motion capture system (Oxford Metrics, Oxford, UK) and post-processed using Vicon Nexus software. Marker trajectories and path data from D-Flow software (Motek Medical, Netherlands) were analyzed in MATLAB (MathWorks, Natick, MA).

We only analyzed the two 3-minute experimental trials for each condition. Marker trajectory data were first low-pass filtered (4th-order Butterworth, cutoff: 10 Hz), then interpolated to 600 Hz (using Matlab function ‘interp1’ with piecewise cubic splines) to ensure accurate stepping-event detection (63). Heel strikes were determined using a marker-based algorithm (64). For consistency, we analyzed the first *N* = 250 steps of each trial.

### Data Analysis

#### Head Pitch Angle

We calculated head pitch angle to quantify participants’ head orientation, as this approximates where people directed their gaze as they walked (65, 66). We defined a local head coordinate system (*x_H_*, *y_H_*, *z_H_*) from the four head markers, taking the origin as the centroid of those four markers. The +*x_H_* axis was directed anterior, the +*y_H_* axis was directed superior, and the +*z_H_* axis was directed laterally toward the right ear, perpendicular to the +*x_H_* and +*y_H_* axes (Fig. 2A). We used a least-squares fitting technique (67) to compute homogeneous transformations between the path and head coordinate systems. Head rotations were calculated from a decomposed ZYX Cardan rotation sequence at each heel strike. We extracted rotation about the *z_H_* axis, or head pitch angle, where downward head pitch is negative (Fig. 2A). We calculated within-trial means, *μ*(*Pitch*), and standard deviations, *σ*(*Pitch*), for each trial.

**Figure 2.**
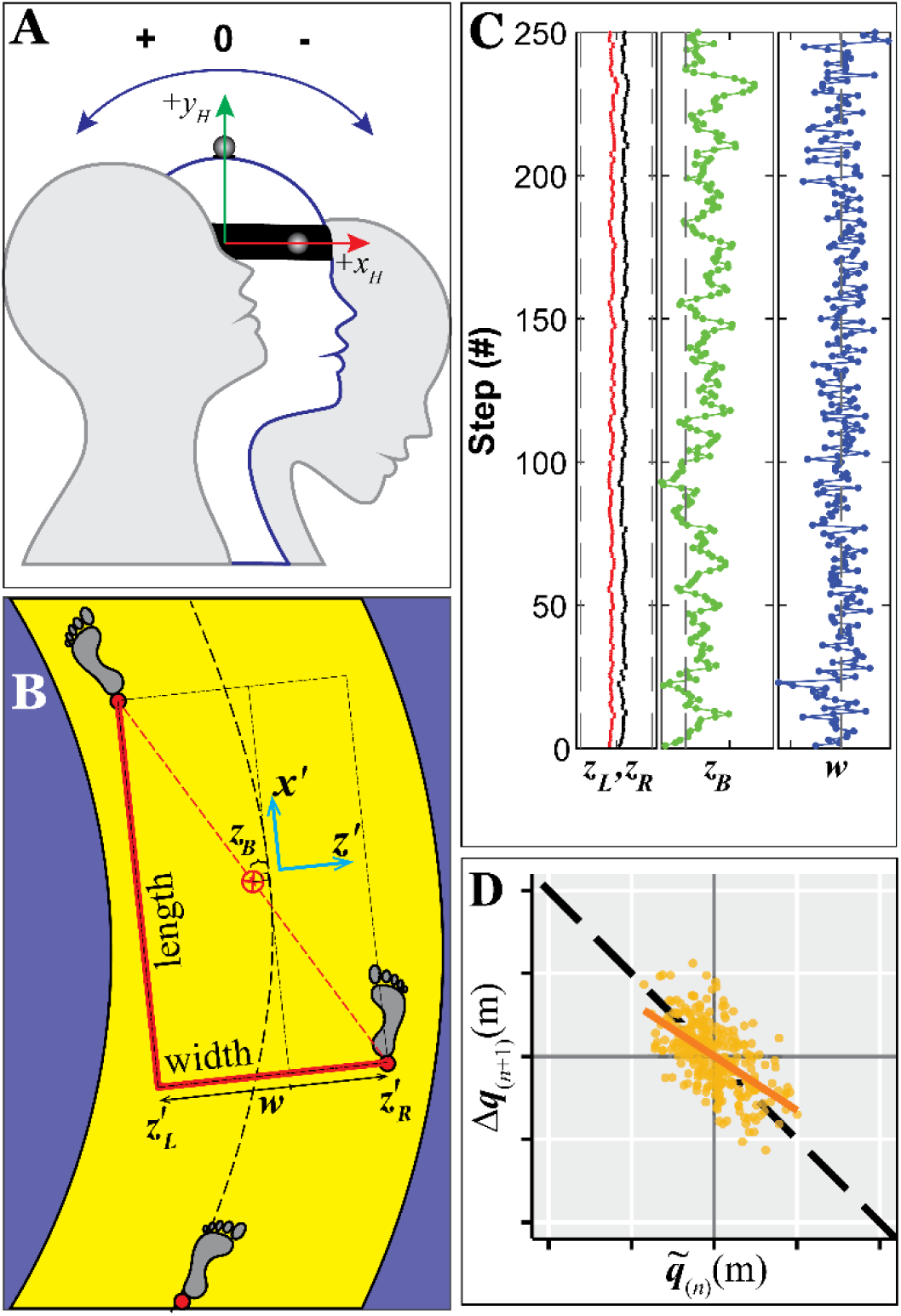
Data Analysis Methods. **A)** Head pitch angle quantifies the sagittal plane rotation of the participant’s head. In the local head coordinate system, the +*x_H_* axis points forward, +*y_H_* axis upward, and +*z_H_* axis (axis of rotation) points outward, perpendicular to the page. Upward head movement creates positive pitch, while downward head movement creates negative pitch. **B)** At each step, we rotate the lab-fixed global coordinate system, [*x,z*], to define a path-aligned local coordinate system, [*xʹ,zʹ*], in which the *xʹ* axis is tangent to the centerline of the path closest to the midpoint between the two feet (6). The locally lateral positions of the left and right feet {*zʹ_L_*, *zʹ_R_*} were then used to compute lateral body position, *z_B_* = ½(*zʹ_L_*+ *zʹ_R_*), and step width, *w* = *zʹ_R_*−*zʹ_L_*, relative to the path. **C)** Example time series data of 250 consecutive steps of left (*z_L_*; red) and right (*z_R_*; black) foot placements, body positions (*z_B_*; green) and step widths (*w*; blue) for a representative trial from a typical participant. In the {*z_L_*; *z_R_*} plot (left), black vertical dashed lines indicate the lateral edges of the treadmill (±0.6 m). In the *z_B_* plot (center), the black vertical dashed line indicates the center of the path (i.e., *z_B_* = 0). In the *w* plot (right), the black vertical dashed line indicates the mean step width. **D)** Example direct control plot used to quantify how people correct errors in *q* ∈ {*z*_B_, *w*} from step-to-step: deviations from the mean on a given step, 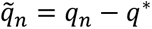, are plotted (yellow markers) against the corrections made on the subsequent step, Δ*q*_(*n*+1)_ = *q*_(*n*+1)_ - *q*_*n*_. The solid yellow diagonal line indicates the error correction slope, *M*(*q*), computed via linear least-squares. The black dashed line indicates “perfect” error correction, i.e.: *M*(*q*) = −1.

We first conducted a mixed-effects analysis of variance (ANOVA) with repeated measures to test the effects of Path (STR vs. HIF) on *μ*(*Pitch*) and *σ*(*Pitch*). To satisfy normality assumptions for *σ*(*Pitch*) we first log-transformed these data. Here, we pooled data across Environment Richness and Path Contrast. In the ANOVA, we treated Path Frequency as a fixed factor, participant as a random factor, and the eight trials (i.e., 2 trials for each of 4 conditions) served as repeated measures.

Subsequently, for each Path separately (STR and HIF), we applied a two-factor (Environment Richness × Path Contrast) mixed-effects ANOVA with repeated measures to test for differences between conditions for each *μ*(*Pitch*) and *σ*(*Pitch*). When significant interaction effects were identified, we performed Tukey’s pairwise post-hoc comparisons to test for differences between Environment Richness (Rich, Sparse) for each Path Contrast (High, Low), and between Path Contrast for each Environment Richness. These statistical analyses were conducted using Minitab (Minitab, Inc., State College, PA). We calculated partial-Eta squared 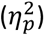 in R (https://www.r-project.org/) to evaluate effect sizes. We interpreted small effects as 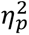 ≥ 0.01, medium effects as 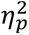 ≥ 0.06, and large effects as 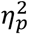 ≥ 0.14 (68).

#### Lateral Stepping Regulation

The Goal Equivalent Manifolds (GEMs) (49) framework allows us to quantify how individuals regulate their lateral stepping across successive steps while walking (19). Using the GEM approach, we determined whether people adapt their lateral stepping regulation in response to changes in perceptual salience on the different Paths presented (STR and HIF).

On a winding path, one’s direction of motion changes at each new step. We therefore transformed our data into path-based local coordinates (69, 70) (Fig. 2B). At each step, we aligned this path-based local coordinate system to be tangent to the point on the centerline of the path closest to the midpoint between the left and right foot placements for that step (Fig. 2B) (69). The locally lateral (*z*-axis) motion was then taken to be perpendicular to the path at that point on the path. Lateral placements of the left and right feet, {*z_Ln_, z_Rn_*}, for each consecutive step *n* ∈ {1,⋯,*N*} were defined as the lateral positions of respective heel markers relative to the path center at each heel strike.

Humans do not regulate {*z_Ln_, z_Rn_*} directly (19). Rather, {*z_Ln_, z_Rn_*} serve as end effectors that enact regulation of step width, *w_n_* = *z_Rn_* − *z_Ln_*, and absolute lateral position relative to their walking path, *z_Bn_* = ½(*z_Ln_* + *z_Rn_*) (Fig. 2B). Foot placements, {*z_Ln_, z_Rn_*}, are thus mathematically related to these goal states, {*z_Bn_, w_n_*}, as (19):

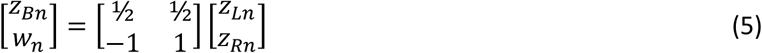

We computed *z_Bn_* and *w_n_* for all *N* = 250 steps throughout each trial (Fig. 2C). For each trial performed by each participant under each condition, we first calculated within-trial means (*μ*) and standard deviations (*σ*) of each *z_B_* and *w* time series.

We then performed linear error correction analyses (47, 71) to quantify the extent to which participants corrected step-to-step deviations in *z_B_* and *w* on subsequent steps (Fig. 2D). We presume people try to maintain (relative to their path) some desired value, *q*\*, for each *q*∈{*z_B_*, *w*}. On any given step, *n*, they will exhibit some small deviation from *q*\*, 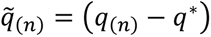. How much they correct those deviations on the subsequent step is then just Δ*q*_(*n*+1)_ = *q*_(*n*+1)_ – *q*_(*n*)_. We directly quantified the extent to which 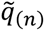 deviations were corrected by Δ*q*_(*n*+1)_ (47, 71). For each *q* ∈{*z_B_*, *w*} time series, we plotted the corrections (Δ*q*_(*n*+1)_) against their corresponding deviations 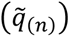 and computed linear slopes, *M*(*q*), from these plots (Fig. 2D). Slopes equal to −1 indicate deviations that are completely corrected at the next step. Conversely, slopes approaching 0 indicate deviations that go uncorrected.

Our previous experiment established that people alter their stepping regulation between walking on straight (STR) and winding (HIF) paths (47, 62). Therefore, for each Path Frequency (STR, HIF), we applied the same two-factor (Environment Richness × Path Contrast) mixed-effects analyses of variance (ANOVA) as described above for each dependent measure: *μ*(*z_B_*), *μ*(*w*), *σ*(*z_B_*), *σ*(*w*), *M*(*z_B_*) and *M*(*w*). To satisfy normality assumptions for *σ*(*z_B_*), and *σ*(*w*), we first log-transformed these data. As warranted, we again performed relevant Tukey’s pairwise post-hoc comparisons as described above.

## RESULTS

### Head Pitch Angle

Across visual conditions, participants adopted overall mean head pitch angles (Fig. 3A), *μ*(Pitch), that averaged −12.35° (IQR: [−20.52°, −3.92°]) on straight (STR) paths, but decreased to −20.12° (IQR: [−25.97°, −13.54°]) on winding (HIF) paths. This decrease was highly significant (F = 227.0; p < 2×10^-16^) with a very large effect size (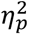 = 0.35). Variability of their head pitch angles (Fig. 3B), *σ*(Pitch), averaged 6.05° (IQR: [3.53°, 8.35°]) on STR paths, but decreased to 3.65° (IQR: [2.58°, 4.32°]) on HIF paths. This decrease was also highly significant (F = 239.6; p < 2×10^-16^) with a very large effect size (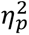 = 0.36). Thus, across variations in visual contexts, participants tilted their heads down substantially more (↓ *μ*(Pitch); Fig. 3A) and maintained this downward head pitch more consistently (↓ *σ*(Pitch); Fig. 3B) when walking on the winding (HIF) paths.

**Figure 3.**
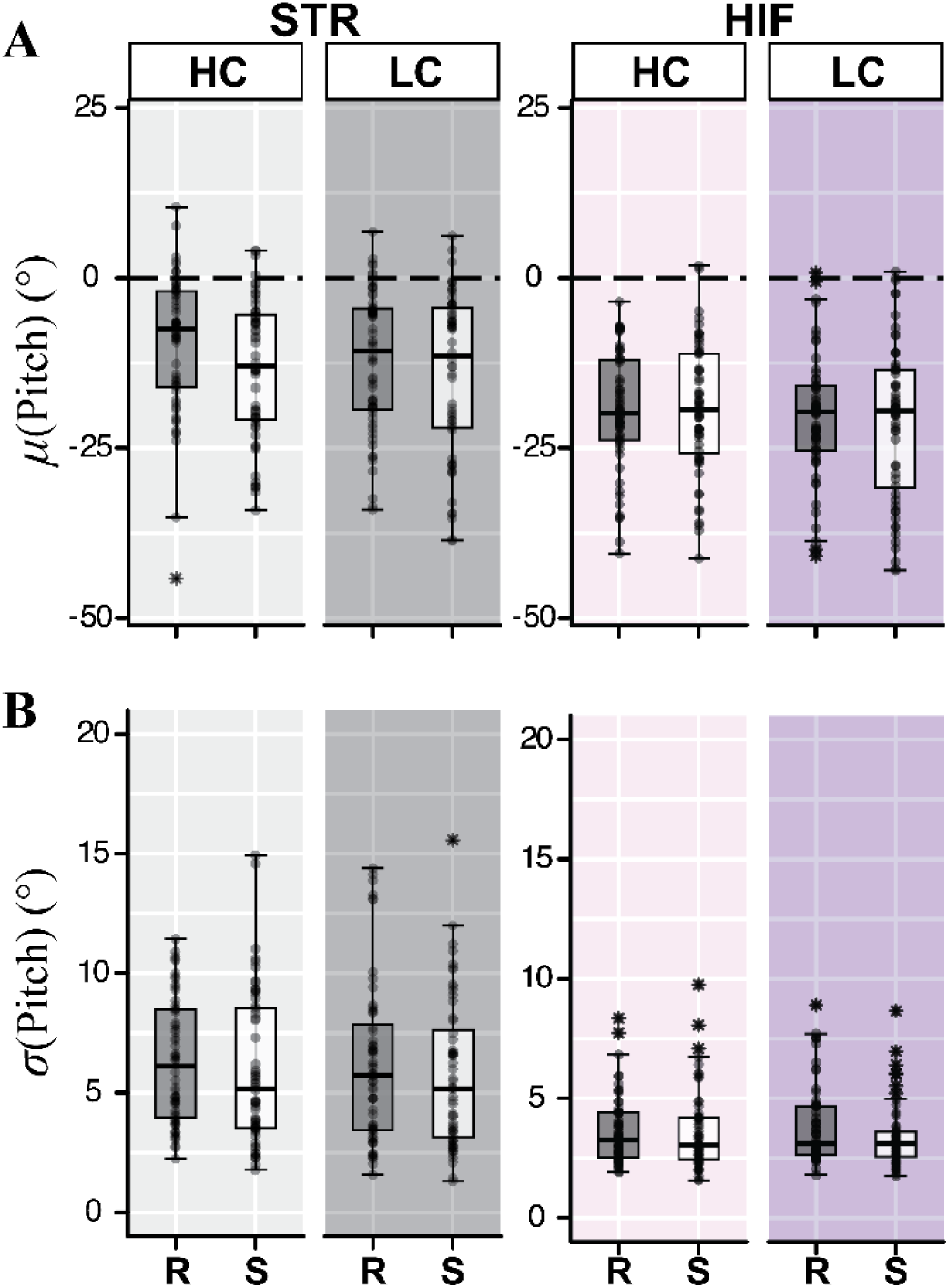
Head Pitch Results. **A)** Within-trial means (*μ*) of head Pitch angle for each environment [Rich (R) and Sparse (S)] and path color contrast [High (HC) and Low (LC)] combination while walking on each path: STR (left) and HIF (right). **B)** Corresponding within-trial standard deviations (*σ*) of head Pitch angle. Box plots show the medians, 1^st^ and 3^rd^ quartiles, and whiskers extending to 1.5 × interquartile range. Values beyond this range are shown as individual asterisks. The overlaid markers are individual data points for each of two trials from each participant. Participants showed significant reductions in *μ*(Pitch) and *σ*(Pitch) while navigating winding paths (p < 0.001). On straight paths (STR), *μ*(Pitch) and *σ*(Pitch) decreased in Sparse environments (p ≤ 0.028) and *μ*(Pitch) decreased on Low Contrast paths (p = 0.03). On winding paths, *μ*(Pitch) decreased on Low Contrast paths (p = 0.021), and *σ*(Pitch) decreased in Sparse environments (p = 0.011). Results of statistical comparisons are in Table 2.

**Table 2.** Head Pitch Statistical Results. Results for ANOVA comparisons between Environment Richness (Rich, Sparse) and Path Contrast (High, Low) for the data shown in Fig. 3, including: means (*μ*) and variability (*σ*) for head Pitch Angle for both straight (STR) and winding (HIF) paths.

| Data In | Variable | Environment | Contrast | Environment<br>× Contrast |
| --- | --- | --- | --- | --- |
| Fig. 3A: | $\mu(\text{Pitch})$<br>STR | $F_{(1,193)} = 27.73$<br>**$p < 0.001$**<br>$\eta_p^2 = 0.13$ | $F_{(1,193)} = 9.09$<br>**$p = 0.003$**<br>$\eta_p^2 = 0.04$ | $F_{(1,193)} = 3.03$<br>$p = 0.083$<br>$\eta_p^2 = 0.02$ |
| | $\mu(\text{Pitch})$<br>HIF | $F_{(1,193)} = 0.10$<br>$p = 0.757$<br>$\eta_p^2 < 0.01$ | $F_{(1,193)} = 5.46$<br>**$p = 0.021$**<br>$\eta_p^2 = 0.03$ | $F_{(1,193)} = 0.90$<br>$p = 0.345$<br>$\eta_p^2 < 0.01$ |
| Fig. 3B: | $\ln(\sigma(\text{Pitch}))$<br>STR | $F_{(1,193)} = 4.89$<br>**$p = 0.028$**<br>$\eta_p^2 = 0.02$ | $F_{(1,193)} = 1.80$<br>$p = 0.182$<br>$\eta_p^2 < 0.01$ | $F_{(1,193)} = 0.07$<br>$p = 0.798$<br>$\eta_p^2 < 0.01$ |
| | $\ln(\sigma(\text{Pitch}))$<br>HIF | $F_{(1,193)} = 6.55$<br>**$p = 0.011$**<br>$\eta_p^2 = 0.03$ | $F_{(1,193)} = 0.20$<br>$p = 0.657$<br>$\eta_p^2 < 0.01$ | $F_{(1,193)} = 2.20$<br>$p = 0.140$<br>$\eta_p^2 = 0.01$ |

ANOVA results (F-statistics, p-values, and $\eta_p^2$ ) are provided for main effects of Environment, Contrast, and Environment × Contrast. Significant differences are indicated in bold.

When walking on the STR paths (Fig. 3, left), introducing a Sparse environment led participants to slightly decrease both their *μ*(Pitch) (p < 0.001) and *σ*(Pitch) (p = 0.028), whereas introducing a Low Contrast path led participants to slightly decrease their *μ*(Pitch) (p = 0.03). When walking on the HIF paths (Fig. 3, right), introducing a Sparse environment led participants to slightly decrease their *σ*(Pitch) (p = 0.011), whereas introducing a Low Contrast path led participants to slightly decrease their *μ*(Pitch) (p = 0.021). However, although these changes were statistically significant, nearly all of them were associated with “small” effect sizes 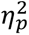 ≤ 0.04, with only one comparison, the decreased *μ*(Pitch) in response to the Sparse environment on STR paths, reaching a “medium” effect size (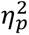 = 0.13; Table 2) (68). Thus, on each path (STR and HIF) separately, changing either the peripheral environment or path contrast led participants to make further small adjustments to their head pitch angles (Fig. 3).

### Lateral Stepping on Straight Paths

Across all visual conditions, participants exhibited similar values (Fig. 4A) for both average lateral positions, *μ*(*z_B_*), and average step widths, *μ*(*w*), when walking on straight (STR) paths (p ≥ 0.068; 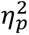 ≤ 0.02; Table 3).

**Figure 4.**
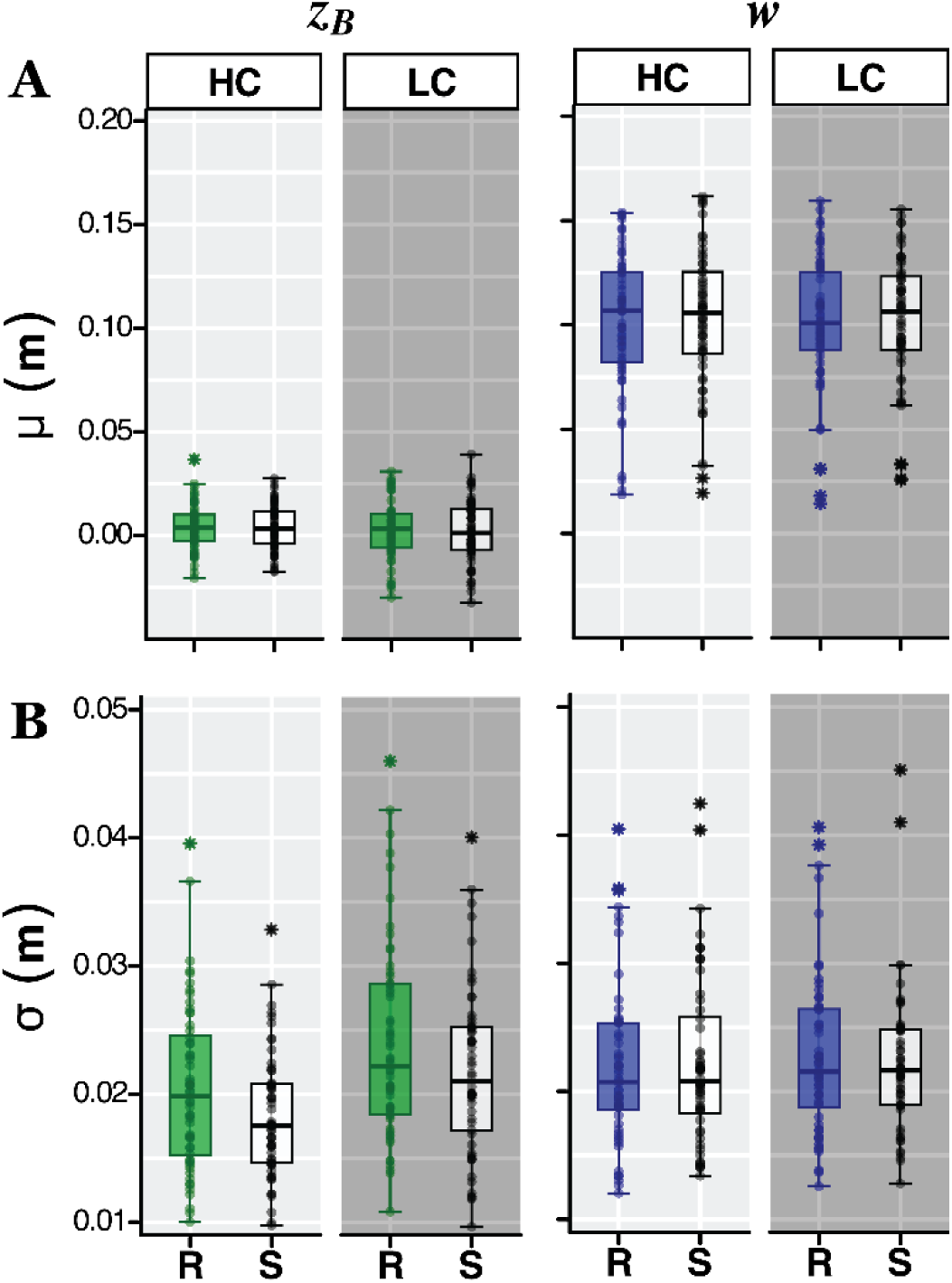
Straight (STR) Path Stepping Results. **A)** Within-trial means (*μ*) for lateral body position, *z_B_* (left), and step width, *w* (right), for each environment [Rich (R) and Sparse (S)] and path color contrast [High (HC) and Low (LC)] combination. **B)** Corresponding within-trial standard deviations (*σ*) for *z_B_* and *w*. Data are plotted in the same manner as in Fig. 3. Variability in lateral position (*σ*(*z_B_*)) decreased in Sparse environments (p < 0.001) but increased on Low Contrast paths (p < 0.001). Conversely, step width variability (*σ*(*w*)) remained consistent across both Environment (p = 0.841) and Path Contrast (p = 0.108). Results of all statistical comparisons are in Table 3.

Introducing a *Sparse environment* led to participants exhibiting slightly less lateral position variability (*σ*(*z_B_*); Fig. 4B; p < 0.001; 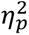 = 0.07; Table 3), and slightly tighter step-to-step corrections of *z_B_* deviations (M(*z_B_*) → −1.0; Fig. 5B; p < 0.001; 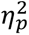 = 0.11; Table 3). Both changes were associated with “medium” effect sizes 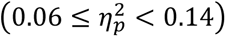 (68).

**Figure 5.**
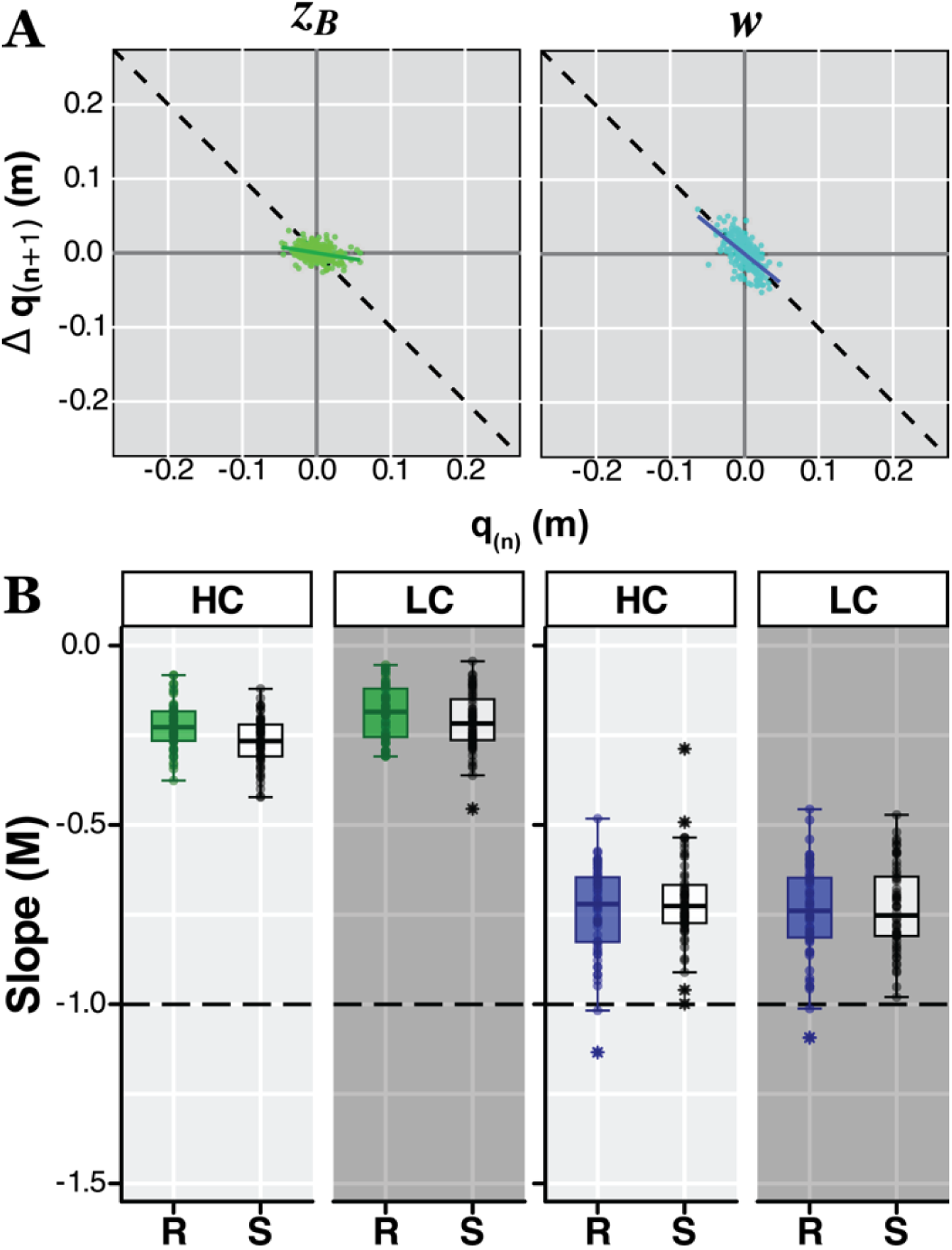
Straight (STR) Path Error-Correction Results. **A)** Representative direct control plots of 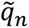 vs. Δ*q*_(n+1)_ (see Fig. 2D) from a typical participant walking on a straight path for lateral body position (*q* ≡ *z*_B_; left) and step width (*q* ≡ *w*; right). **B)** Group data for error correction slopes for lateral body positions, *M*(*z_B_*) (left), and step widths, *M*(*w*) (right), for each environment [Rich (R) and Sparse (S)] and path color contrast [High (HC) and Low (LC)] combination. Data are plotted in the same manner as in Figs. 3-4. Horizontal dashed lines at slope *M* = −1 indicate perfect error correction: i.e., values above this line reflect *under*-correction, whereas values below this line reflect *over*-correction.

**Table 3.** Straight (STR) Path Stepping Statistical Results. Results for ANOVA comparisons between Environment Richness (Rich, Sparse) and Path Contrast (High, Low) for data shown in Figs. 4-5: means (*μ*) and variability (*σ*) for lateral position (*z_B_*) and step width (*w*), and direct control slopes (*M*) for *z_B_* and *w*.

| Data In | Variable | Environment | Contrast | Environment<br>× Contrast |
| --- | --- | --- | --- | --- |
| Fig. 4A: | $\mu(z_B)$ | $F_{(1,193)} = 0.33$<br>$p = 0.567$<br>$\eta_p^2 < 0.01$ | $F_{(1,193)} = 2.89$<br>$p = 0.091$<br>$\eta_p^2 = 0.01$ | $F_{(1,193)} = 0.07$<br>$p = 0.791$<br>$\eta_p^2 < 0.01$ |
| | $\mu(w)$ | $F_{(1,193)} = 3.37$<br>$p = 0.068$<br>$\eta_p^2 = 0.02$ | $F_{(1,193)} = 0.53$<br>$p = 0.468$<br>$\eta_p^2 < 0.01$ | $F_{(1,193)} = 0.03$<br>$p = 0.863$<br>$\eta_p^2 < 0.01$ |
| Fig. 4B: | $\ln(\sigma(z_B))$ | $F_{(1,193)} = 15.05$<br>**$p < 0.001$**<br>$\eta_p^2 = 0.07$ | $F_{(1,193)} = 42.83$<br>**$p < 0.001$**<br>$\eta_p^2 = 0.18$ | $F_{(1,193)} = 0.00$<br>$p = 0.949$<br>$\eta_p^2 < 0.01$ |
| | $\ln(\sigma(w))$ | $F_{(1,193)} = 0.04$<br>$p = 0.841$<br>$\eta_p^2 < 0.01$ | $F_{(1,193)} = 2.61$<br>$p = 0.108$<br>$\eta_p^2 = 0.01$ | $F_{(1,193)} = 1.03$<br>$p = 0.311$<br>$\eta_p^2 < 0.01$ |
| Fig. 5B: | $M(z_B)$ | $F_{(1,193)} = 22.68$<br>**$p < 0.001$**<br>$\eta_p^2 = 0.11$ | $F_{(1,193)} = 42.58$<br>**$p < 0.001$**<br>$\eta_p^2 = 0.18$ | $F_{(1,193)} = 1.06$<br>$p = 0.305$<br>$\eta_p^2 < 0.01$ |
| | $M(w)$ | $F_{(1,193)} = 1.93$<br>$p = 0.167$<br>$\eta_p^2 < 0.01$ | $F_{(1,193)} = 0.89$<br>$p = 0.347$<br>$\eta_p^2 < 0.01$ | $F_{(1,193)} = 0.74$<br>$p = 0.392$<br>$\eta_p^2 < 0.01$ |

ANOVA results (F-statistics, p-values, and $\eta_p^2$ ) are provided for main effects of Environment, Contrast, and Environment × Contrast. Significant differences are indicated in bold.

Conversely, introducing a *Low Contrast path* led to participants exhibiting greater lateral position variability (*σ*(*z_B_*); Fig. 4B; p < 0.001; 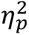 = 0.18; Table 3), whereas they corrected step-to-step *z_B_* deviations slightly less (M(*z_B_*) → 0.0; Fig. 5B; p < 0.001; 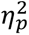 = 0.18; Table 3). These changes were associated with “large” effect sizes (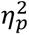 > 0.14) (68).

Across visual conditions, step-to-step dynamics of step widths (*w*) remained remarkably consistent. Participants exhibited no significant differences in mean step widths (*μ*(*w*); Fig. 4A), variability of step widths (*σ*(*w*); Fig. 4B), or step-to-step error-correction of step widths (*M*(*w*); Fig. 5B) for variations in either peripheral Environment or path Contrast (all p ≥ 0.068; all 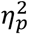 ≤ 0.02; Table 3). Thus, when walking on straight (STR) paths, participants made minor adjustments to *z_B_*-regulation in Sparse environments, somewhat greater adjustments to *z_B_*-regulation on Low Contrast paths, but maintained consistent *w*-regulations across these visual manipulations.

Participants corrected *z_B_* deviations more in Sparse environments (p < 0.001) and less on Low Contrast paths (p < 0.001). Conversely, they corrected *w* deviations similarly across both Environment and Path Contrast (p ≥ 0.167). Results of all statistical comparisons are in Table 3.

### Lateral Stepping on Winding Paths

On the winding (HIF) paths, participants exhibited substantially different stepping dynamics than they did on the straight (STR) paths. They took narrower steps (*μ*(*w*); Fig. 6A vs. Fig. 4A), substantially more variable steps (*σ*(*z_B_*) & *σ*(*w*); Fig. 6B vs. Fig. 4B) (see also Fig. 7A vs. Fig. 5A), and exhibited much stronger step-to-step error correction (↓ *M*) for both lateral position and step width (*M*(*z_B_*) & *M*(*w*); Fig. 7B vs. Fig. 5B). These large differences were consistent with findings from our prior work on similar winding paths (47), and corroborate the increased stepping control challenges of walking on winding paths.

**Figure 6.**
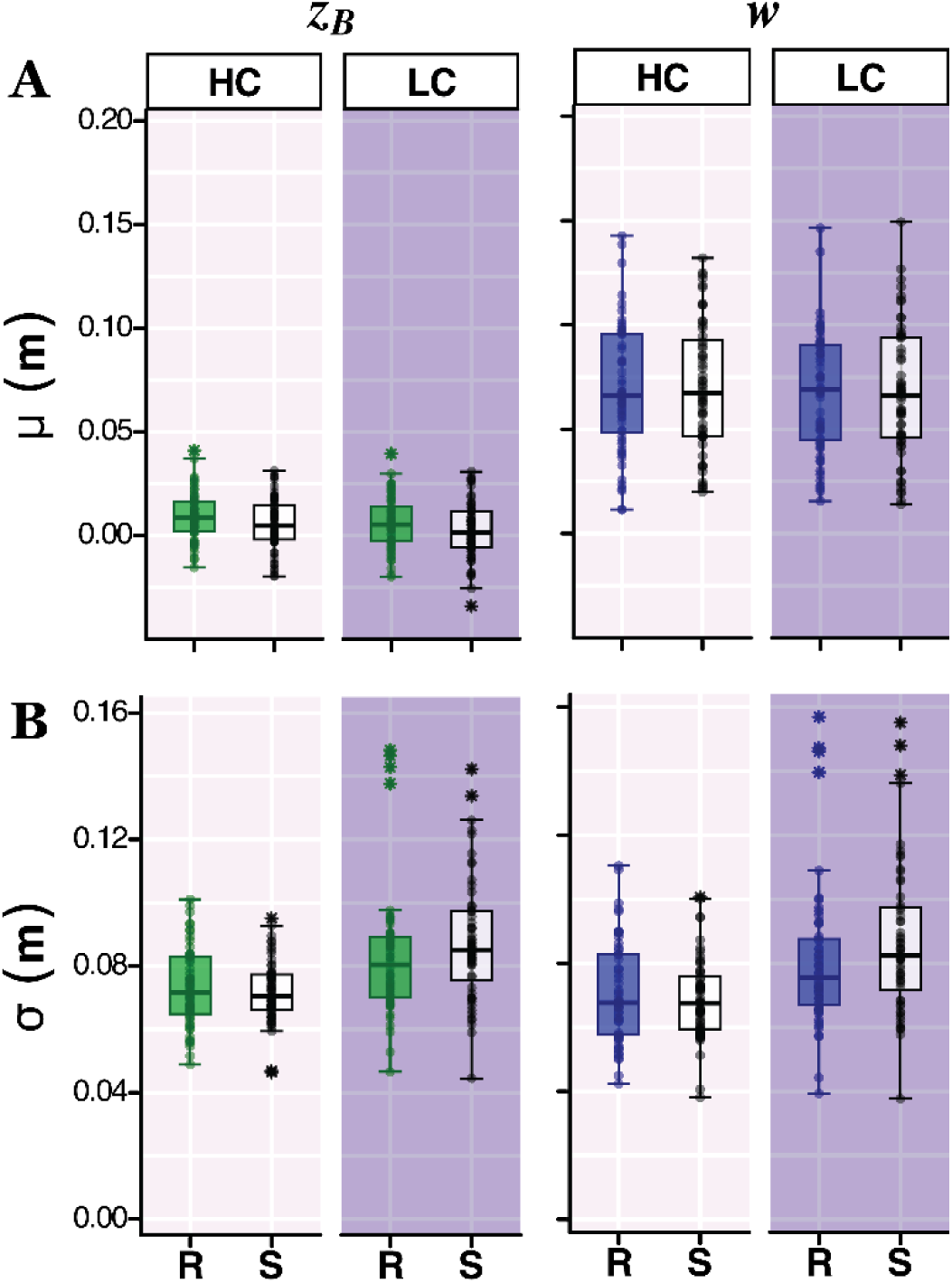
Winding (HIF) Path Stepping Results. **A)** Within-trial means (*μ*) for lateral body position, *z_B_* (left), and step width, *w* (right), for each environment [Rich (R) and Sparse (S)] and path color contrast [High (HC) and Low (LC)] combination. **B)** Corresponding within-trial standard deviations (*σ*) for *z_B_* and *w*. Data are plotted in the same manner as in Figs. 3-5. Mean lateral position (*μ*(*z_B_*)) decreased in Sparse environments (p < 0.001) and on Low Contrast paths (p ≤ 0.002). Variability of both lateral position (*σ*(*z_B_*)), and step width (*σ*(*w*)) increased with lower path contrast overall (p < 0.001). *σ*(*w*) further increased on Low Contrast paths in Sparse environments (p = 0.033). Results of all statistical comparisons are in Table 4.

**Figure 7.**
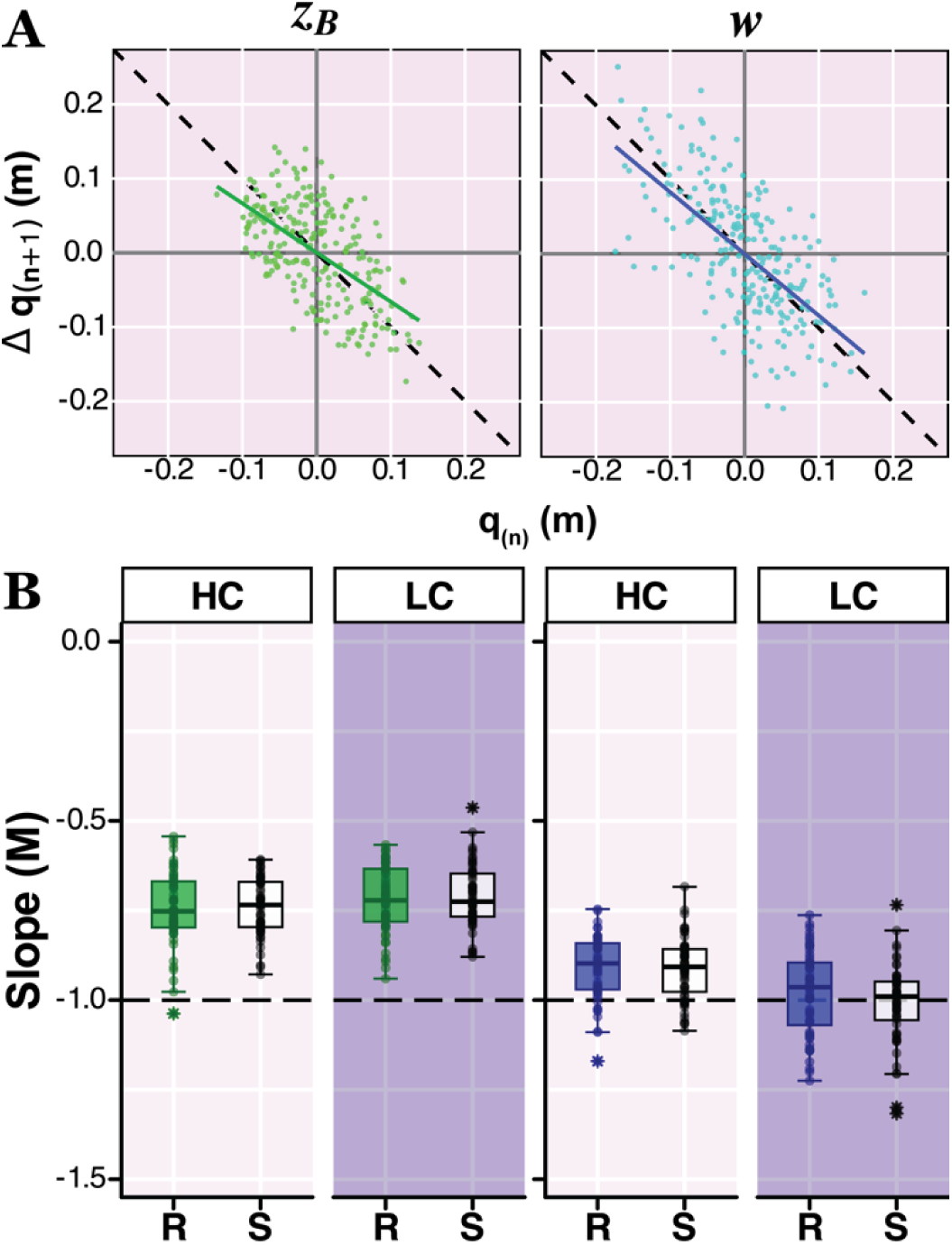
Winding (HIF) Path Error-Correction Results. **A)** Representative direct control plots from a typical participant for lateral body position (*z*_B_; left) and step width (*w*; right). **B)** Group data for error correction slopes for lateral body positions, *M*(*z_B_*) (left), and step widths, *M*(*w*) (right), for each environment [Rich (R) and Sparse (S)] and path color contrast [High (HC) and Low (LC)] combination. Data are plotted in the same manner as in Fig. 5. Participants corrected *z_B_* deviations (*M*(*z_B_*)) less (p < 0.001) and *w* deviations (*M*(*w*)) more (p < 0.001) on Low Contrast paths. Results of all statistical comparisons are in Table 4.

Across all visual conditions, participants exhibited consistent mean step widths (*μ*(*w*); Fig. 6A; p ≥ 0.052; 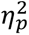 ≤ 0.02; Table 4).

**Table 4.** Winding (HIF) Path Stepping Statistical Results. Results for ANOVA comparisons between Environment Richness (Rich, Sparse) and Path Contrast (High, Low) for data shown in Figs. 6-7: means (*μ*) and variability (*σ*) for lateral position (*z_B_*) and step width (*w*), and direct control slopes (*M*) for *z_B_* and *w*.

| Data In | Variable | Environment | Contrast | Environment<br>× Contrast | Tukey's<br>(Environment Effects) | Tukey's<br>(Contrast Effects) |
| --- | --- | --- | --- | --- | --- | --- |
| Fig. 6A: | $\mu(z_B)$ | $F_{(1,193)} = 14.26$<br>**$p < 0.001$**<br>$\eta_p^2 = 0.07$ | $F_{(1,193)} = 9.93$<br>**$p = 0.002$**<br>$\eta_p^2 = 0.05$ | $F_{(1,193)} = 0.05$<br>$p = 0.829$<br>$\eta_p^2 < 0.01$ | N/A | N/A |
| | $\mu(w)$ | $F_{(1,193)} = 0.38$<br>$p = 0.540$<br>$\eta_p^2 < 0.01$ | $F_{(1,193)} = 3.83$<br>$p = 0.052$<br>$\eta_p^2 = 0.02$ | $F_{(1,193)} = 0.58$<br>$p = 0.447$<br>$\eta_p^2 < 0.01$ | N/A | N/A |
| Fig. 6B: | $\ln(\sigma(z_B))$ | $F_{(1,193)} = 1.82$<br>$p = 0.178$<br>$\eta_p^2 < 0.01$ | $F_{(1,193)} = 73.97$<br>**$p^* < 0.001$**<br>$\eta_p^2 = 0.28$ | $F_{(1,193)} = 4.40$<br>**$p = 0.037$**<br>$\eta_p^2 = 0.02$ | High Contrast:<br>Rich-Sparse: $p = 0.952$<br>Low Contrast:<br>Rich-Sparse: $p = 0.073$ | Rich:<br>**HC-LC: $p < 0.001$**<br>Sparse:<br>**HC-LC: $p < 0.001$** |
| | $\ln(\sigma(w))$ | $F_{(1,193)} = 2.04$<br>$p = 0.155$<br>$\eta_p^2 = 0.01$ | $F_{(1,193)} = 78.50$<br>**$p^* < 0.001$**<br>$\eta_p^2 = 0.29$ | $F_{(1,193)} = 6.01$<br>**$p = 0.015$**<br>$\eta_p^2 = 0.03$ | High Contrast:<br>Rich-Sparse: $p = 0.887$<br>Low Contrast:<br>**Rich-Sparse: $p = 0.033$** | Rich:<br>**HC-LC: $p < 0.001$**<br>Sparse:<br>**HC-LC: $p < 0.001$** |
| Fig. 7B: | $M(z_B)$ | $F_{(1,193)} = 1.74$<br>$p = 0.188$<br>$\eta_p^2 < 0.01$ | $F_{(1,193)} = 25.81$<br>**$p < 0.001$**<br>$\eta_p^2 = 0.12$ | $F_{(1,193)} = 0.15$<br>$p = 0.695$<br>$\eta_p^2 < 0.01$ | N/A | N/A |
| | $M(w)$ | $F_{(1,193)} = 2.55$<br>$p = 0.112$<br>$\eta_p^2 = 0.01$ | $F_{(1,193)} = 67.74$<br>**$p < 0.001$**<br>$\eta_p^2 = 0.26$ | $F_{(1,193)} = 1.57$<br>$p = 0.212$<br>$\eta_p^2 < 0.01$ | N/A | N/A |

ANOVA results (F-statistics, p-values, and $\eta_p^2$ ) are provided for main effects of Environment, Contrast, and Environment × Contrast. Significant differences are indicated in bold. \*In cases where Environment × Contrast interactions were significant, we considered main effects results to be unreliable and potentially misleading. We drew conclusions instead from the Tukey's pairwise comparisons, as these account for the effects of the interactions.

Introducing a *Sparse environment* led to small decreases in mean lateral position (*μ*(*z_B_*); Fig. 6A; p < 0.001; 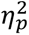 = 0.07; Table 4) and increased step width variability (*σ*(*w*); Fig. 6B), but only on the Low Contrast paths (p = 0.033; Table 4). However, environment richness had no effect on the extent to which participants corrected either *z_B_* deviations (M(*z_B_*); Fig. 7B; p = 0.188; 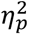 < 0.001; Table 4) or *w* deviations (M(*w*); Fig. 7B; p = 0.112; 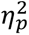 = 0.01; Table 4).

Conversely, introducing a *Low Contrast path* led to participants exhibiting much greater variability of both lateral position, *σ*(*z_B_*), and step width, *σ*(*w*) (Fig. 4B; p < 0.001; Table 4). Participants corrected *z_B_*-deviations (lateral position) less (M(*z_B_*) → 0.0; Fig. 7B; p < 0.001; 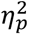 = 0.12) and corrected *w* deviations more (M(*w*) → −1.0; Fig. 7B; p < 0.001; 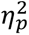 = 0.26) on these Low Contrast paths. These changes in step-to-step error correction were each strongly significant (p < 0.001) and associated with “medium” 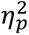 = 0.12 and “large” 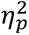 = 0.26 effect sizes, respectively (Table 4) (68). Thus, when walking on winding (HIF) paths, participants exhibited far more substantial changes in stepping regulation in response to changes in path contrast (Low vs. High) than in response to changes in environmental richness (Sparse vs. Rich).

## DISCUSSION

Walking is a fundamental activity that plays a crucial role in our daily lives, enabling us to navigate our environments and engage with the world around us (1, 2, 4, 5). However, the ability to walk is significantly influenced by sensory input, particularly vision (20, 22, 23, 31, 35, 45). Visual information helps us to perceive our surroundings, assess potential paths, and make informed decisions about our movements (3, 21, 24, 39). Our study examined how the quality of that visual information, specifically perceptual salience, affects the embodied stepping decisions people make while walking along specified paths. We manipulated both environment richness and path color contrast and asked participants to walk along both straight and winding paths. We applied the GEM framework and tested how these variations in visual information influenced lateral stepping regulation.

Our guiding hypothesis, that walking involves continuous decision-making action-perception processes (9, 10, 21), gave rise to three plausible outcomes. Our experimental design held contextual salience constant for each walking path (straight or winding) and independently manipulated the perceptual salience of both the peripheral environment and walking paths to directly assess each of the three plausible outcomes predicted. By testing these complementary predictions, our research sought to elucidate how perception shapes goal-directed motor behavior, bridging theoretical frameworks (19, 43, 46) with practical navigation challenges in real-world-like settings.

The first predicted outcome was that, since the biomechanical requirements of each task (i.e., the contextual salience) were held constant and participants had full vision to be able to adequately see each path, stepping regulation would remain consistent across the different visual contexts presented. This outcome was largely supported in so far as differences between the two walking tasks (straight vs. winding paths) elicited far greater changes to stepping regulation than did either visual manipulation within each walking task. Nonetheless, additional differences due to the visual manipulations, independent of each walking task, indicate that merely “seeing” one’s path is not sufficient, but that the quality and quantity of the visual information available also informs ongoing stepping decisions.

The second predicted outcome was that, if gaze is used primarily to enhance *task-relevant* visual information (3, 38) and reduce visual uncertainty (23, 24, 45), then diminishing the visual contrast of the paths, but not the surrounding peripheral environment, would degrade participants’ stepping control. This outcome was supported in so far as changing the path contrast elicited more and greater changes to stepping regulation than did changing the peripheral environment. But it was not the case that manipulating the surrounding peripheral environment led to *no* changes in stepping control.

The third predicted outcome was that peripheral vision would also contribute to governing locomotor behavior (22, 35, 36, 54), such that decreasing the perceptual salience of either the peripheral environment or the paths would induce participants to change their stepping behavior. This prediction was also supported, although to a lesser extent than the effects of manipulating the path visual saliency or the contextual salience of the tasks themselves (i.e., straight vs. winding paths). Taken together, these findings demonstrate how contextual salience, perceptual salience, and cognitive salience (46) interact during a naturalistic walking task (e.g., as in (9), etc.), down even to the level of choosing stepping movements during ongoing walking (18, 19, 48).

Overall, participants showed a notable reduction in average head pitch angle and variability while navigating winding paths compared to straight paths (Fig. 3). This was consistent with them using gaze to obtain, and decrease uncertainty of, *task-relevant* information (23, 24, 38) about their walking paths. Humans readily adjust their gaze and head position in response to their environment (65, 66), and in particular to surface complexity (72). As surface complexity increases, people lower their heads to direct their gaze toward the ground to gather critical visual information to ensure stable gait (72–74) and accurate foot placement (3, 21, 72, 75). Winding paths pose more complex navigational challenges than straight paths (76), requiring continuous adjustments to foot placement (47, 77, 78). The lower head pitch and lower head pitch variability on winding paths thus aligns with previous research and indicates that dependence on visual information shifts toward ground-directed feedback for more accurate foot placement. This influence of path shape on head pitch modulation suggests that path curvature plays a significant role in how people interact with visual cues while walking. Based on this, we expected variations in environmental richness and path contrast to affect walkers differently depending on path shape. Our within-path findings support this expectation, as these visual manipulations led to distinct changes in head position and stepping behavior between straight and winding paths.

On straight paths, young, unimpaired walkers often direct their gaze farther ahead (3, 31, 33, 34). However, while walking in sparse environments, participants adopted a head tilt that was pitched further down (medium effect) and less variable (small effect) (Fig. 3). This adjustment may be attributed to the minimal visual elements in the Sparse environment, which limited information from both peripheral vision and optic flow. As a result, walkers placed greater emphasis on the path itself rather than on peripheral environmental cues (24, 38). Moreover, changes in optic flow can induce gait adaptations including shorter steps, and greater variability in step width, step length (79), stride length, time, and speed, fore aft position on the treadmill (80), and walking velocity (81). Here, people adjusted their lateral foot placements to modify their lateral body position, but not step width, on straight paths (Figs. 4-5). In these Sparse environments, participants corrected deviations in lateral body position more (medium effect) (Fig. 5) resulting in decreased lateral position variability (medium effect) (Fig. 4). This compensation for reduced spatial orientation in Sparse environments due to its lack of positional cues, allowed people to maintain a more consistent position on the path.

When the path’s boundaries were less distinguishable on straight, low color contrast paths, participants tilted their heads further downward (small effect) (Fig. 3A) but maintained variability similar to the high contrast paths (Fig. 3B). This small decrease in head pitch suggests increased attention directed towards the path itself. However, this did not translate into tighter control over their lateral positioning. Indeed, participants corrected deviations in lateral body position less (large effect) (Fig. 5), which in turn resulted in increased lateral body position variability (large effect) (Fig. 4). This suggests that despite slightly increasing attention toward the path, individuals still struggled to maintain consistent position when the path’s boundaries were less distinguishable. Alternatively, that participants did not exert tighter control over lateral position may instead reflect that they chose to prioritize maintaining strong step width regulation (Fig. 5) (19).

On winding paths, changes in path contrast more strongly influenced stepping adaptations than did changes in the richness of the surrounding environment. Walking in sparse environments only decreased Head Pitch variability (small effect) (Fig. 3). All other adaptations were influenced by the contrast in path colors. This follows from the overall lower average head pitch people adopted while navigating winding paths in general (Fig. 3), indicating a heightened focus on the path itself. On low color contrast winding paths, participants tilted their heads further down (small effect) (Fig. 3). Although lower head pitch may indicate a greater reliance on ground-directed feedback (24, 38) for accurate stepping, both lateral body position and step width were more variable while walking along low color contrast paths (Fig. 6). Challenges of precise foot placement were exacerbated while walking in sparse environments, leading to increased step width variability (Fig. 6) due to limited visual cues from both the path and environment. Furthermore, participants corrected deviations in lateral body position (Fig. 7) less on low color contrast paths (medium effect), consistent with the increased lateral position variability (Fig. 6). However, step widths were also generally more variable, but this is likely due to participants overcorrecting for deviations in step width (large effect) (Fig. 7). This suggests that individuals prioritized adjustments in step width over lateral position corrections in response to degraded path information on winding paths.

The observed variations in stepping regulation highlight the brain’s capacity to integrate visual information and adapt motor outputs in real-time (8, 14, 38). This supports the predictions that perceptual salience does affect embodied decisions during walking, independent of the biomechanical demands of the task. Thus, walking is not merely a mechanical task, but an embodied decision-making process (8, 51) influenced by both contextual and perceptual factors (22, 24, 28, 38). While the quality of available visual information does affect how individuals regulate their lateral stepping, the impact is not especially strong. Rather, contextual salience, specifically path shape, had a more substantial impact on lateral stepping (47). This suggests that young individuals can – for the most part – readily adapt their gaze (here, head pitch) to acquire additional visual information needed to enact appropriate stepping regulation when navigating less perceptually salient contexts.

Understanding how the salience of visual information impacts walking behaviors can inform the design of environments, particularly for individuals with mobility challenges or those navigating unfamiliar spaces. Enhancing environmental richness and optimizing color contrast in public spaces could facilitate safer and more effective navigation, ultimately improving the quality of life for individuals across varying mobility levels.

In conclusion, our study highlights specifically how perceptual salience interacts with contextual salience in shaping walking behavior and demonstrates the adaptability of individuals in response to changing visual contexts. The findings contribute to a growing body of literature that emphasizes the dynamic interplay between perception and action in goal-directed locomotion. By connecting theoretical concepts and practical applications, we can deepen our understanding of walking behaviors and inform strategies to improve navigation in complex environments.

## DATA AVAILABILITY

Source data can be found on DataDryad: https://doi.org/10.5061/dryad.7sqv9s56j

## ACKNOWLEDGMENTS

The authors thank Carlie J. Hernjac and Sarah E. Overby for their assistance with conducting the experiments, data collection, and discussions on interpreting the findings.

Present address of A.C.R.: University of Utah, HPER East, 260 S 1850 E, Salt Lake City, UT 84112.

## AUTHOR CONTRIBUTIONS

A.C.R., T.S., J.P.C., and J.B.D. conceived and designed research; A.C.R. performed experiments, A.C.R. analyzed data; A.C.R., T.S., J.P.C., and J.B.D. interpreted results of experiments, A.C.R. prepared figures and drafted manuscript; A.C.R., T.S., J.P.C., and J.B.D. edited and revised manuscript; A.C.R., T.S., J.P.C., and J.B.D. approved final version of manuscript.

